# Neurodegenerative disease associated pathways in brain of the triple transgenic Alzheimer’s model are reversed in vivo following two weeks peripheral administration of fasudil

**DOI:** 10.1101/2022.09.30.510301

**Authors:** Richard Killick, Christina Elliott, Elena Ribe, Martin Broadstock, Clive Ballard, Dag Aarsland, Gareth Williams

## Abstract

The pan ROCK inhibitor fasudil acts as a vasodilator and has been used as a medication post cerebral stroke for the past 27 years in Japan and China. More recently, on the basis of the involvement of ROCK inhibition on synaptic function, neuronal survival and processes associated with neuroinflammation, it has been suggested that the drug may be repurposed for neurodegenerative diseases. Indeed, fasudil has demonstrated preclinical efficacy in many neurodegenerative disease models.

To facilitate an understanding of the wider biological processes at play due to ROCK inhibition in the context of neurodegeneration we performed a global gene expression analysis on the brains of Alzheimer’s disease model mice treated with fasudil via peripheral i.p injection. Our results show that fasudil tends to drive gene expression in a reverse sense to that seen in postmortem neurodegenerative disease brains. The results are most striking in terms of pathway enrichment analysis where pathways regulated in Alzheimer’s disease and by fasudil treatment are overwhelmingly regulated in opposite directions.

Thus, our results bolster the repurposing potential of fasudil by demonstrating an anti-neurodegenerative phenotype in a disease context and highlight the potential of in vivo transcriptional profiling of drug activity.

## Introduction

Drug repurposing is the application of an existing therapeutic, with a known safety profile and prescription data, to a new condition. This strategy is particularly relevant to neurodegenerative conditions that have so far proved refractory to traditional target based novel entity development[1]. Repurposing offers a quick route to the clinic following the identification of a novel target for the given disease, where one can adopt an off the shelf approach benefiting from the extensive drug development data and post approval patient histories. In this context, inhibition of the Rho-associated coiled-coil containing protein kinases (ROCKs), a target for vasodilation post ischemia for which there is a clinically approved drug, has emerged as a possible strategy in the treatment of neurodegenerative diseases (NDDs).

Fasudil (HA-1077), a vasodilator, was developed by the Hidaka group during the 80s[2] and initially thought to be a Ca^2+^ channel antagonist [3, 4] with particularly potent effects on the vasculature supplying the brain[2, 4]. However, following the identification of ROCK it very soon became apparent that the drug acts on blood vessels through the antagonistic effects on ROCK. There are two ROCK paralogues, ROCK1 and ROCK2, and fasudil has very similar IC50s towards each, 730 and 720 nM, respectively [2]. Fasudil gained clinical approval for the treatment of cerebral vasospasm following subarachnoid hemorrhage (SAH) in Japan in 1994 and then in China[5]. Interestingly, fasudil was observed to also improve neuronal function in animal models of SAH[6]. This led Kamei et al., to test fasudil in two human subjects with cerebrovascular dementia with wondering [7]. Given orally at 30 or 60 mg a day for 8 weeks, fasudil prevented wandering and improved memory in these two subjects.

A genomic association study implicated the human WW1C/KIBRA gene in memory function[8]. The observation that the KIBRA is negatively regulated by ROCK led Huentalman et al[9] to test and show that ROCK inhibition, with hydroxyfasudil (the active metabolite of fasudil), can potentiate memory and even reduce age-dependent memory impairment in rats. This was one of the first reports on the preclinical use of fasudil in the West and gained the attention of the popular media, generating headlines such as: “Drug found that could reduce risk of Alzheimer’s disease”, as appeared in the online publication ScienceDaily[10].

In this context, RhoA/ROCK signaling activity is intimately involved in synapse remodeling[11] and retraction[12] and inflammatory responses in many tissues including brain[13]. Like Alzheimer’s disease (AD), many neurodegenerative diseases (NDDs) also feature synapse loss and neuroinflammation[14]. As such fasudil, and a number of other small molecule antagonists of ROCK, have been used in numerous preclinical studies in animal models across many different NDDs[15]. In these studies, fasudil has repeatedly been reported to have beneficial effects[16-19].

To investigate the mechanistic action of fasudil in the context of NDD pathology further, we performed a global transcriptional analysis of brain tissue from fasudil treated 3xTg-AD mice. The triple transgenic model captures the two major AD neuropathologies, senile plaque by 6 months and neurofibrillary tangles by 12 months. At 4 months cognitive impairment is also observed[20]. After peripheral administration of fasudil via i.p. injection regimen over two weeks we found significant gene expression changes in the brains of the treated mice. By comparing our expression profile with data generated from publicly available post-mortem NDD (AD, Parkinson’s disease (PD) and Huntington’s disease (HD)) sample data we show that fasudil tends to drive gene expression in an opposite sense to that in NDD. This effect is more pronounced in the genes down regulated in NDD and emerges most clearly through a pathway enrichment analysis. Our results highlight the potential of transcription profiling to reveal the therapeutic potential of candidate therapeutics in the context of NDD. Specifically, we have shown how expression profiling of fasudil activity at the site of disease pathology reveals its disease ameliorating potential through its driving gene expression in an opposite sense to that found in disease.

## Methods

### Drug treatment

At 18 months of age 3xTg-AD mice were administered 10 mg/kg fasudil (n=7) or PBS vehicle (n=10), twice daily, i.p., for two weeks. Mice were sacrificed and brains removed, hemisected, and fixed or snap frozen as previously described[16]. Rostral cortices were subsequently collected from the snap frozen hemibrains after partial thawing on ice and total RNA extracted by homogenisation directly in TRIzol reagent and isopropanol precipitation, then further purified using RNeasy ® mini kit (Ref: 74104, Qiagen, US).

### Microarray analysis

RNA was converted to cRNA and hybridised to full genome Affymetix Mouse Gene 2.0 ST microarrays, processed and imaged according to the manufacturer’s instructions. The CEL files were processed in the R environment using the Bioconductor oligo package to give RMA normalised expression values[21].

The self-organising map (SOM) analysis[22] was performed on the expression data after it was normalised to *N*(0,1) across samples. Random weights were defined on an 10×10 square lattice and mapping optimisation performed using a Euclidean distance metric. The iteration is as follows:

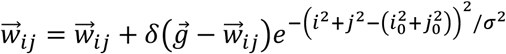

where 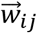 are the SOM weights and the vector refers to the space of samples, 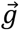 is expression level of the given gene across the samples and the weight closest to 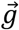 is at (*i*_0_, *j*_0_). The step size *δ* is set to 0.01 and reduced by a factor of 0.99 upon each iteration. The weight neighbourhood extends over the Gaussian standard deviation *σ* and this is initialised at 2 and reduced by a factor of 0.99 upon each iteration.

The weights were then regressed against treatment status or animal sex. A further SOM was performed on the residual expression data with sex variance subtracted.

Expression profiles were defined as the linear fit Z scores for treatment versus control samples. The fasudil samples comprised both male and female mice and the Z scores were generated based on a linear model with sex as a covariate. The probes were mapped to genes with the maximal magnitude Z score selected in cases of alternative probes.

### Meta-analysis transcription profiles

The AD profile was based on the composite Z scores for 21 profiles from 17 series corresponding to post-mortem brain samples that were publicly available on the NCBI GEO data repository[23], as described by us in[24]. The PD composite profile was defined in a similar way to the AD profile and based on the 12 GEO expression series: GSE8397[25], GSE46036[26], GSE20295[27], GSE20164[28], GSE49036[29], GSE7621[30], GSE19587[31], GSE34516[32], GSE20314[28], GSE43490[33], GSE54282[34], GSE24378[28]. This resulted in a composite from 21 distinct PD profiles. The HD profile was based on the GEO series GSE3790[35]. Here, the expression levels were fit with a linear model with age, sex and brain region as covariates.

### Pathway analysis

Pathway analysis was based on the distribution of pathway genes on the full Z score-based expression profiles. Statistical significance was based on an analytical function parametrised by the maximal difference in positive to negative enrichment in the cumulative curve, as described by us previously[36].

### CMAP pathway enrichment profiles

The Connectivity MAP 2.0[37] comprises gene expression change data for the activity of 1,309 drug-like compounds on cancer cell lines. The data was downloaded in the form of ranked probes for the given drug treatments relative to the plate controls. This data was mapped to average probe ranks for each compound resulting in 1,309 profiles. Pathway profiles, defined as the lists of significantly enriched pathways for the expression profiles, were then generated for each compound. The various neurodegenerative disease associated pathway profiles were compared to each of the CMAP pathway profiles and ranked according to the ratio of pathways that are regulated in the same direction to those that are regulated in the opposite direction. The graphs are for 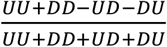, where *U* and *D* correspond to the positive and negative enrichment.

## Results

### Expression changes in the brains of 3xTg-AD mice post fasudil treatment

To assess the global transcriptional effects on the brains of AD model mice post fasudil treatment, we performed a SOM analysis, as described in Methods above. A SOM consists of a matrix of weights each capturing distinct gene expression variation in the experiment. The amount of variation explained by fasudil treatment can be gauged by regressing the weights against treatment status. We find that the SOM segregates into islands representing genes whose expression is up/down regulated by fasudil, see Figure 1. Thus, it appears that fasudil has a significant effect on overall gene expression. A similar analysis with sex points to a more modest expression change, see Figure 1. We further generated a SOM with the effects of sex subtracted with a linear model fit across the samples. Here, the fasudil effects are more striking, see Figure 1.

**Figure 1.**
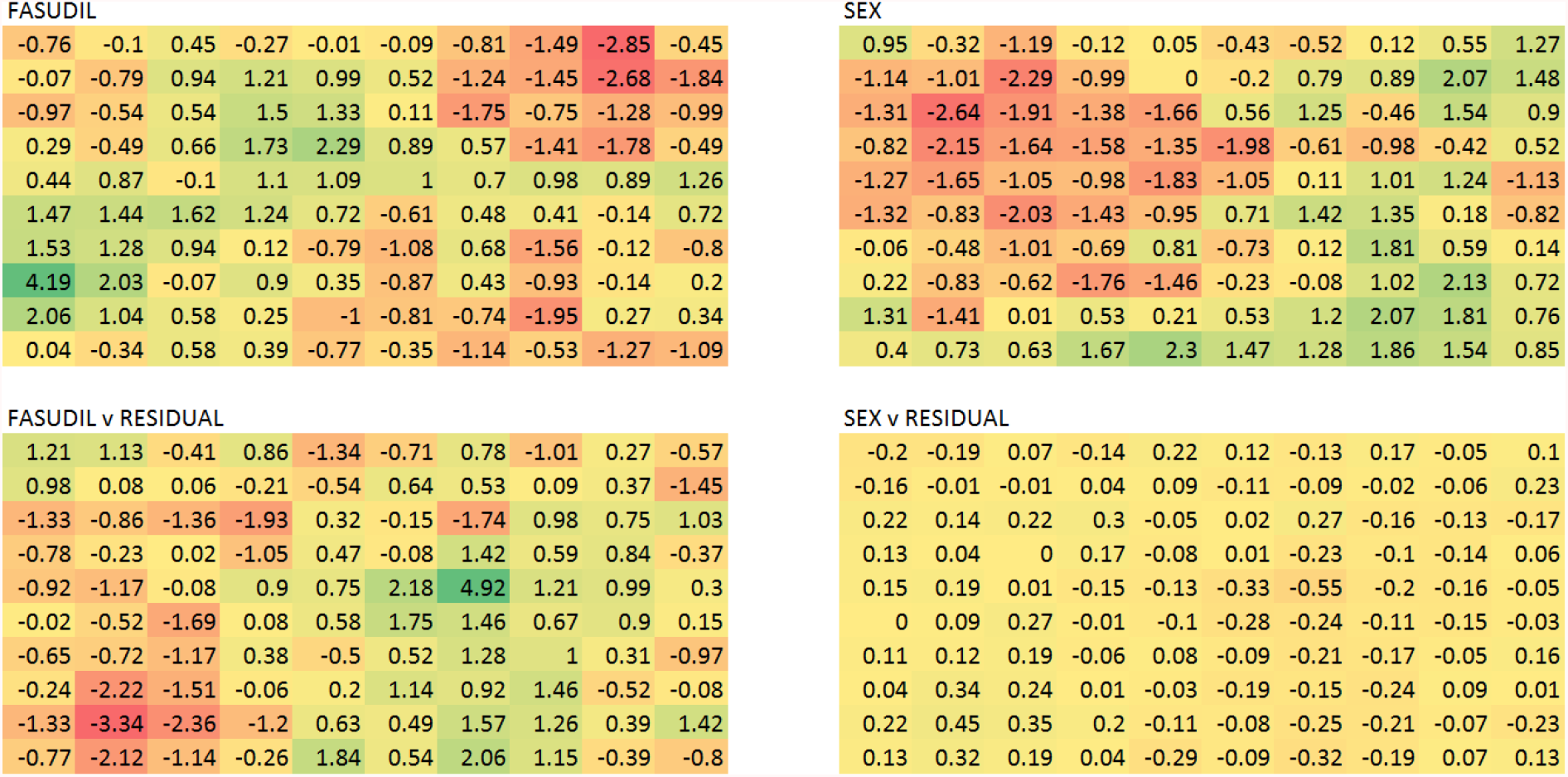
Self-organising maps of the transcriptional data reveal the global effects of fasudil treatment. In the top left the SOM weights are regressed against treatment status and the corresponding Z scores shown. It is clear that there are islands of positive and negative correlation corresponding to genes that are up and down regulated respectively. On the top right the weights are correlated with the mouse sex and show a significant sex effect that is however muted relative to that of fasudil. At the bottom the SOM for the residual expression data resulting from the sex component subtraction is show with regression against treatment and sex on the left and right respectively. As expected, sex no longer shows any significant correlation with the weights and interestingly the fasudil correlations are stronger.

Performing a differential expression analysis, we found that the expression changes driven by fasudil in the brains of the 3xTg-AD mice were relatively modest with only 9 and 1 genes significantly (p < 0.05) altered at the two-fold level in the female animals. In the males there were 3 up and 1 down two-fold gene changes and full profiles controlling for sex only 2 up and one down regulated gene at this level. Restricting analysis to these genes will only furnish limited analysis of biological changes driven by fasudil. We therefore decided to base our analysis on the global expression profile defined as the ranked Z scores corresponding to the gene changes between control and treatment groups. As described in the Methods section we defined a fasudil profile by fitting expression levels to treatment category and sex. The profile then consists of 15,707 genes of which 479 and 414 are up and down regulated at two standard deviations away from the mean respectively.

### Fasudil driven gene expression and neurodegenerative disease

To get an idea regarding the biological systems that are being perturbed by fasudil treatment we performed a pathway analysis along the lines described in detail in our recent paper[36]. Here, the full set of genes in the expression profiles were ranked according to the Z scores of the expression changes across the samples and a pathway enrichment analysis performed.

We find an up regulation of NGF signalling genes (p<1.78E-4) as well as pathways down regulated in AD[24]: PD (KEGG)(p<8.59E-4) and Huntington’s disease (HD) (KEGG) (p<3.59E-3). Oxidative phosphorylation has been shown to be down regulated in AD [Mitochondria dysfunction in the pathogenesis of Alzheimer’s disease: recent advances[38-41] and we find that fasudil up-regulates metabolic pathways: Oxidative Phosphorylation (p<8.17E-4), Voxphos (p<9.93E-3), Mitochondria (p<2.93E-3), Respiratory electron transport (p<3.52E-4). Wnt signalling is reported to be down regulated in AD[42, 43] and we find that fasudil drives this pathway up (p<5.29E-3).

The observation that pathways associated with AD and other neurodegenerative diseases are significantly regulated led us to investigate a possible overlap with transcriptional changes in the AD brain. To this end we generated pathways for a composite AD profile described in[44]. At the level of perturbed genes, we can see that there is indeed a significant anti-correlation with AD and other NDD profiles (PD and HD), as shown in the top ranked gene list and the contingency table at |Z| > 2 threshold, Figure 2 and Table 1. The reversal is particularly evident with those genes down regulated in AD. Of the shared genes at the Z > 2 level with 20 of the top 25 most down regulated genes in AD being up regulated by fasudil treatment. The picture is less clear for the genes up regulated in AD, with 14 in the top 25 down regulated by fasudil treatment.

**Table 1.**
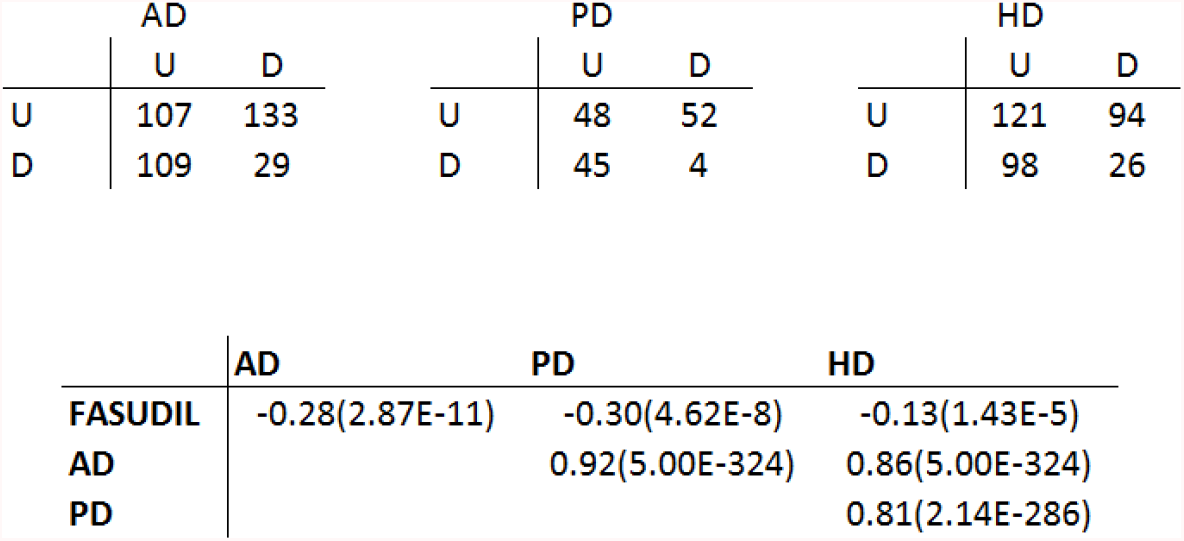
The correlations of the fasudil transcription profile with neurodegenerative disease profiles. The contingency tables for regulated genes at the |*Z*| > 2 level are shown at the top for AD, PD and HD. The corresponding correlation scores, measured by 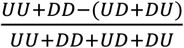 where the *U* and *D* stand for up and down regulated genes, are shown below. The significance by one sided Fisher exact test is shown in brackets. The fasudil profile significantly anti-correlates with the AD, PD and HD profiles. The neurodegenerative disease profiles all show strong positive correlations with each other.

|  | AD |  |  | PD |  |  | HD |  |
| --- | --- | --- | --- | --- | --- | --- | --- | --- |
|  | U | D |  | U | D |  | U | D |
| U | 107 | 133 | U | 48 | 52 | U | 121 | 94 |
| D | 109 | 29 | D | 45 | 4 | D | 98 | 26 |

|  | AD | PD | HD |
| --- | --- | --- | --- |
| FASUDIL | -0.28(2.87E-11) | -0.30(4.62E-8) | -0.13(1.43E-5) |
| AD |  | 0.92(5.00E-324) | 0.86(5.00E-324) |
| PD |  |  | 0.81(2.14E-286) |

**Figure 2.**
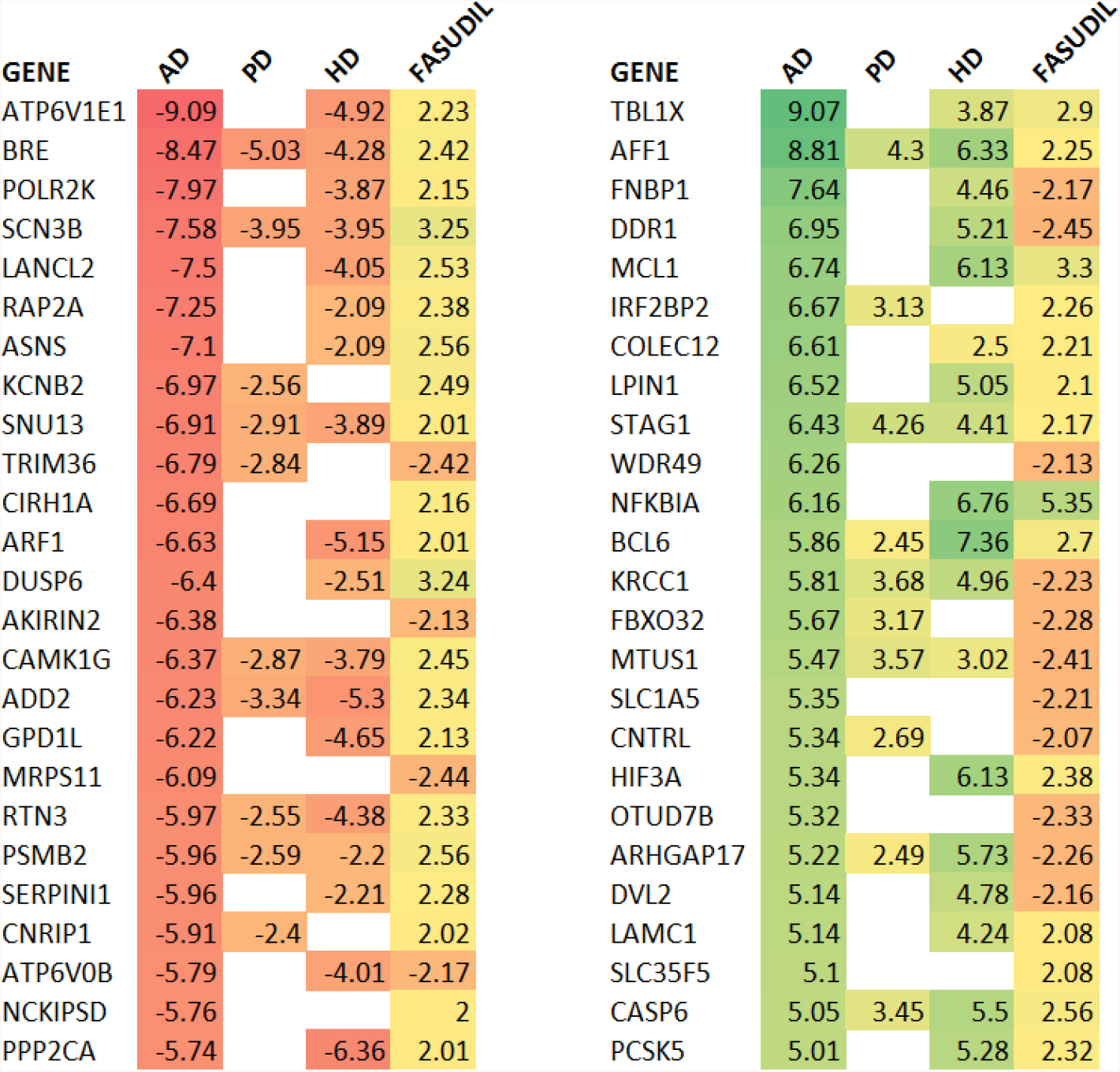
Fasudil driven gene expression changes tend to be the reverse of those in multiple neurodegenerative conditions. The gene expression Z scores are shown for AD, PD and HD together with those for the fasudil treatment. The most down regulated genes in AD are shown at the left and the most up regulated at the right. As expected, there is a high degree of consistency across the other neurodegenerative conditions, PD and HD. Fasudil shows an up regulation of all but four genes of those most down regulated in AD. The reversal is less significant for those genes up regulated in AD with only 14 out of 25 driven down with fasudil.

A pathway enrichment analysis provides an insight into the underlying biology and can facilitate a comparison between expression profiles that is less sensitive to the noisy component in gene expression. In this context it is interesting to observe that the contrast between NDD and fasudil is more clearly delimited through a comparative pathway analysis. An overall picture of the pathways regulated in NDD and by fasudil is given in Figure 3 and the contingency table plus statistics in Table 2. In the pathway set down regulated in AD those that are regulated by fasudil are overwhelmingly driven upwards (all in the top 25). In the top 25 up regulated pathways seen in AD 19 are down regulated by fasudil treatment.

**Table 2.** The correlations of the fasudil treatment regulated pathway set with those perturbed in neurodegenerative disease profiles. The contingency tables for regulated pathways at the |*Z*| > 2 level are shown at the top for AD, PD and HD. The corresponding correlation scores, measured by 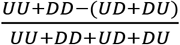 where the *U* and *D* stand for up and down regulated genes, are shown below. The significance by one sided Fisher exact test is shown in brackets. The fasudil profile significantly anti-correlates with the AD, PD and HD profiles. The pathway analysis provides a stronger case for the anti-neurodegenerative potential of fasudil that the gene wise analysis.

|  | AD |  |  | PD |  |  | HD |  |
| --- | --- | --- | --- | --- | --- | --- | --- | --- |
|  | U | D |  | U | D |  | U | D |
| U | 45 | 104 | U | 45 | 66 | U | 73 | 52 |
| D | 74 | 0 | D | 20 | 3 | D | 46 | 6 |

|  | AD | PD | HD |
| --- | --- | --- | --- |
| FASUDIL | -0.60(7.10E-28) | -0.28(3.74E-5) | -0.11(4.78E-5) |
| AD |  | 0.98(5.47E-105) | 0.96(2.22E-122) |
| PD |  |  | 0.99(3.42E-96) |

**Figure 3.**
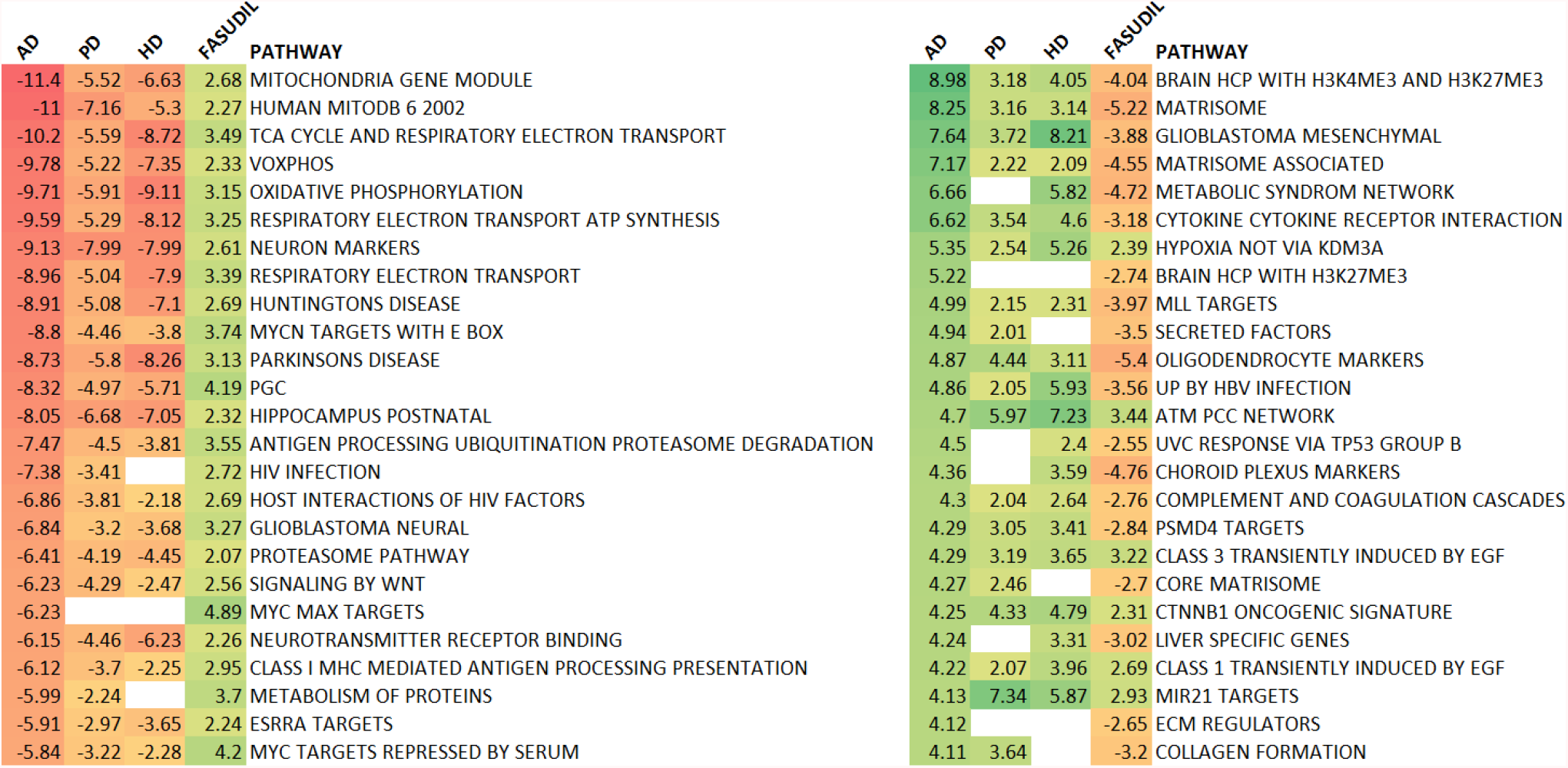
Fasudil regulates pathways in an opposite sense to that seen in multiple neurodegenerative conditions. The pathway enrichment Z scores are shown for AD, PD and HD together with those for the fasudil treatment. The most down regulated pathways in AD are shown at the left and the most up regulated at the right. As expected, there is a high degree of consistency across the other neurodegenerative conditions, PD and HD. Fasudil shows an up regulation of all the pathways most down regulated in AD and a down regulation of 76% of the most up regulated pathways in AD.

### Fasudil driven transcription in vitro

We performed pathway enrichment analyses on all the CMAP compounds as described in Methods. The pathway profiles of the CMAP drugs enabled us to do an extensive comparison with the AD pathway profile. We find that fasudil again anti-correlates with the AD pathway profile and is in the top 9% of anti-correlating profile, see Figure 4. A similar result is seen with PD and HD, Figure 4. Interestingly, a similar analysis based on gene wise comparisons fails to show any anti-correlation between the CMAP fasudil profile and that of AD, which can be performed through the www.spied.org.uk web tool[45]. The latter observation and the relatively weak NDD anti-correlation at the pathway level speaks to the importance of assessing compound activity in a disease relevant context as the therapeutic potential of fasudil would have been missed with an analysis of cell culture based compound expression profile data base searches, such as provided by CMAP and LINCS[46].

**Figure 4.**
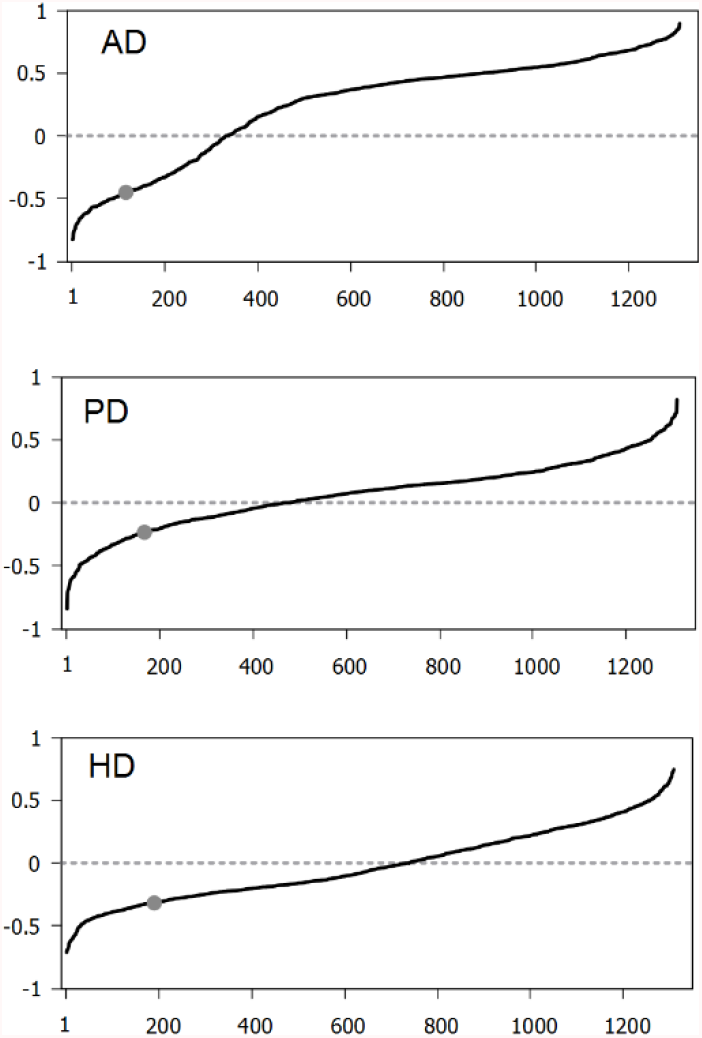
Neurodegenerative disease pathway profile correlations with CMAP pathway profiles. The CMAP profiles are ranked based on the correlation score 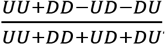, were *U* and *D* correspond to the number of shared pathways regulated upwards and downwards respectively. Fasudil has a relatively high anti-correlation with the NDD profiles. In particular, the contingency table for AD is 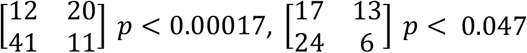 for PD and 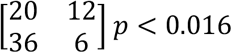 for HD.

## Discussion

Several groups, including our own, have reported observations which demonstrate that the administration of the pan ROCK inhibitor, fasudil, to animal models of AD is beneficial [16, 17, 47-49]. Here, for the first time, we examined the transcriptomic effects of fasudil in the frontal cortex of an AD mouse model, the triple transgenic model, following administration of fasudil by i.p. injection twice daily at 10 mg/kg for two weeks.

An initial global analysis of the microarray gene expression data through a self-organising map reveals a significant effect of fasudil in the frontal cortexes of treated animals. With a view to uncovering the potential AD reversing effects of fasudil we compared its differential expression profile with composite expression profiles derived from various neurodegenerative diseases. Interestingly, we find that there is a significant anti-correlation between the fasudil profile and that of AD, PD and HD. This is particularly evident in the genes down regulated in the neurodegenerative conditions, with 80% of the most down regulated genes in AD being up regulated by fasudil. Differential expression analysis is complicated by the contribution of noise and the large gene to sample ratio. One way of smoothing noise is to perform a pathway enrichment analysis, as this is an analysis based on sets of genes where we expect noise effects to cancel. To this end we performed a pathway enrichment analysis on the fasudil driven expression profile and the composite neurodegenerative disease profiles. Comparing the pathways significantly regulated in disease and drug treatment we find a clear reversal particularly in the pathways down regulated in disease. Here, of the top 25 down regulated pathways in AD that are regulated by fasudil all are driven upwards by the drug.

The potential of pathway enrichment analysis in drug repurposing is highlighted by our analysis of deposited data on drug driven expression profiles in cell lines. In particular, our composite neurodegenerative disease pathway sets show a relatively high anti-correlation with those of fasudil in the CMAP database. For example, fasudil is in the top 9% of hits for AD ordered by anti-correlation. This is in contrast with a gene wise analysis, where there is no significant anti-correlation between the neurodegenerative profiles and fasudil in the same CMAP database. However, the pathway analysis at the *in vitro* level reveals a much weaker connection between fasudil and disease than seen *in vivo*. It will be interesting to see whether the anti-neurodegenerative phenotype of fasudil is revealed in the *in vivo* context because we are looking at responses in the appropriate tissue or whether these disease reversing effects will only be manifest in a disease model.

Our own pervious work has shown fasudil is able to protect synapses in primary rat cortical neuronal cultures against synaptotoxic amounts of oligomeric forms of Aβ1-42 (Aβo)[17], to protect Aβo-driven memory impairment in an acute Aβ rat model[16], and to lower levels of both soluble Aβ and insoluble, plaque-like deposits, of Aβ in multiple brain regions of the same animals examined here. This novel analysis of the transcriptional effects of peripherally delivered fasudil on brain tissue not only demonstrate the drug brain permeable and strengthens our contention that it will be of benefit for treating AD but also supports the contentions of others that fasudil could also be of benefit for treating PD and other synucleinopathies[50] and HD[51, 52]. The data we present here, supporting the use of fasudil for AD, PD and HD, is further supported by an earlier proteomic-based analysis of fasudil which pinpointed the same three neurodegenerative diseases [53].

